# A thermally stable protein nanoparticle that stimulates long lasting humoral immune response

**DOI:** 10.1101/2022.12.30.522068

**Authors:** Ten-Tsao Wong, Gunn-Guang Liou, Ming-Chung Kan

## Abstract

Thermally stable vaccine platform is considered to be the missing piece of vaccine technology. In this article, we are reporting the development of a novel protein nanoparticle and evaluating its ability in withstanding extended high temperature incubation and stimulating long lasting humoral immune response. This protein nanoparticle is assembled from a fusion protein composed of an amphipathic helical peptide derived from M2 protein of H5N1 influenza virus (AH3) and a super folder green fluorescent protein(sfGFP). The proposed structure of this protein nanoparticle is modeled according to transmission electronic microscope (TEM) images of protein nanoparticle. From this proposed protein model, we have generated a mutant with two gain-of-function mutations that function synergistically on particle stability. Protein nanoparticle assembled from this gain-of-function mutant is able to remove a hydrophobic patch from the surface of protein nanoparticle. This gain-of-function mutant also contributes to higher thermostability of protein nanoparticle and stimulates long lasting humoral immune response after single immunization. This protein nanoparticle shows increasing particle stability in higher temperature and higher salt concentration. This novel protein nanoparticle may serve as a thermal-stable platform for vaccine development.

## Introduction

Thermostability of a vaccine is considered an important characteristic that is essential to fulfill Global vaccination initiative. Current vaccine logistic practice demands continuous cold chain environment from manufacturer to remote clinics to maintain vaccine efficacy. It is estimated that there are 775 million people living out of electrical grid (IEA, 2022) and beyond the reach of vaccine cold chain and vaccination which leads to millions of children died of vaccine preventable diseases each year (Kumar et al., 2022). To fully cover the population beyond electrical grid, a vaccine has to be able to withstand temperature up to 40°C for two months, the duration of vaccine stored in health post (Karp et al., 2015).

Subunit vaccine is a safe alternative to traditional inactivated or attenuated vaccines, but its efficacy is often hindered by low antigenicity of recombinant protein. Different approaches have been utilized to resolve this issue, among them, virus like particle (VLP) and self-assembled protein nanoparticle (SAPN) are considered the best platforms for subunit vaccine development (Lopez-Sagaseta et al., 2016). The size of VLPs are ranged between 20-200 nm that facilitates both draining efficiently to lymph node and also uptake by antigen presenting cells like dendritic cell and macrophage (Ahsan et al., 2002). The other benefit of VLP based vaccine is the induction of B cell receptor clustering when presenting repetitive antigen to B cell, a function that can activate antibody class-switch and somatic hypermutation in a T cell dependent mechanism (Bachmann and Jennings, 2010). Not only the virus like particle can be used for vaccine directly, heterologous antigen can be presented in the particle surface through genetic fusion (Lu et al., 2015). The Universal flu vaccine candidate, ectopic M2 domain (M2e), has been genetically fused with Hepatitis B core antigen(HBc) and assembled into a nanoparticle that provides full protection to heterologous flu strains (Ravin et al., 2015). VLP purified from bacterial expressed capsid protein often encloses bacterial RNA that is required for particle stability (Manolova et al., 2008). Although the enclosed bacterial RNA serves as adjuvant during immunization, the labile nature of RNA also contributes to the low stability of VLP. A SAPN that is assembled strictly from protein components may provide better overall vaccine stability. The SAPN based on iron transporting protein, ferritin, has been chosen as vaccine carrier for being constituted strictly by protein subunit (Johnston et al., 2022; Kim et al., 2022). It has good thermostability when exposing to high temperature but the long term storage data for ferritin based vaccine has not been reported yet.

The green fluorescent protein is a member of fluorescent protein family that are structurally conserved and emits fluorescent light from a chromophore when excited by photons of shorter wavelength (Tsien, 1998). The shared features of fluorescent proteins including a sturdy barrel shaped structure constituted by 11 β-sheets and an enclosed chromophore that emits fluorescent light when excited (Dove et al., 2001). The function of the barrel shell is to provide a well organized chemical environment to ensure the maturation of chromophore and protects it from hostile elements (Cody et al., 1993). So it is conceivable that the protein sequences among fluorescent protein family members in the barrel shell are highly variable and fluorescent proteins possess desirable biophysical properties can be selected using directed evolution (Close et al., 2014; Cormack et al., 1996; Crameri et al., 1996; Kiss et al., 2009; Tsien, 1998). The applications of fluorescent protein have been expanded into multiple areas beyond live imaging which includes serving as biological sensors (Pavoor et al., 2009; Wang et al., 2014), or detectors for protein-protein interaction or protein folding (Cabantous et al., 2005; Romei and Boxer, 2019).

Amphipathic α-helical peptide (AHP) forms hydrophilic and hydrophobic faces when folded and is often identified in proteins related to phospholipid membrane interaction. The N-terminal amphipathic helical peptide is required for membrane anchorage of Hepatitis C virus NS3 protein and the protease function of the NS3/NS4a complex (He et al., 2012; Horner et al., 2012). Several anti-microbial peptides also possess amphipathic properties and function by forming membrane pores or causing membrane disruption (Tossi et al., 2000). The amphipathic α-helical of type A influenza virus M2 protein is required for M2 protein anchorage and induces membrane curvature required for virus budding (Roberts et al., 2013; Rossman et al., 2010). Here in this report, we are describing the identification and design of a protein nanoparticle assembled from a fusion protein composing an amphipathic α-helical peptide (AH3) from M2 protein of type A influenza strain H5N1 and a sfGFP. We have built a protein structure model based on TEM images using Ascalaph Designer. The model suggests AH3 serves as a polymerization module and it induces fusion protein assembly into a corn-on-a-cob structure. The sturdy barrel structure of sfGFP was explored as antigen presenting module for peptide antigen presentation. From the protein model, we have generated a gain-of-function mutant that provides higher stability to the protein nanoparticle. We shows biophysical and immunological evidences that the protein nanoparticle assembled from fusion protein containing gain-of-function mutant is able to withstand extended high temperature exposure and stimulates long lasting antibody response to an inserted antigen in single immunization without adjuvant.

## Materials and Methods

### Peptide information, expression and purification of recombinant protein

The AH3 peptide sequence is DRLFFKCIYRRLKYGLKRG. The sequence of peptide for anti-hM2e antibody titer ELISA is SLLTEVETPIRNEWGSRSNGSSDC. The peptide inserted in the insertion site of sfGFP is SLLTEVETPIRNEWGSRSN-GSSDSSGGSLLTEVETPIRNEWGSRSNGSSD. The protein expression vectors encoding target proteins were transformed into E coli competent cells using a heat shock transformation. Colonies of transformed bacteria with the desired vector were scraped from plate and inoculated in LB culture with an antibiotic. The bacterial culture was then growing exponentiallyd to OD600 between 0.5∼0.7 before cooling down on an ice bath and protein expression was induced by 1mM IPTG at 20 °C for 14-16 hours with 250 rpm shaking. After protein induction, bacteria were harvested by 5000 rpm centrifugation in a Sorvall SLC3000 rotor for 10 minutes. Bacterial pellet from 400 ml LB culture was re-suspended in 40 ml lysis buffer contains 10 mM Imidazole in 1XGF buffer (20 mM Na(PO_4_) pH7.4 and 300mM NaCl) for sonication. For Ni-NTA resin purification: 10 mM Imidazol was added in 1XGF buffer for bacteria lysis (Lysis buffer), 20 mM Imidazol was added in 1XGF buffer for column wash (Wash buffer), 500 mM Imidazol was added in 1XGF buffer for protein elution (Elution buffer). Bacteria were lysed using an ultrasonic sonicator (Misonix 3000) at 10 second on/20 second off cycles for 5 minutes at output level 5 in icy water.

Insoluble cell debris was removed by centrifugation in 10000 rpm for 10 minutes using a Sorval SS34 rotor at 4 °C. Soluble fraction containing the target protein was then used for purification by Ni-NTA resin as described in the user manual or been used for sedimentation ultracentrifugation. Purified proteins were stored in elution buffer in cold room before further processing. Before immunization or thermostability test, protein buffer was changed into 0.5x GF buffer using Sephadex-25 resin (GE, PD-10 and NAP-5).

### Biophysical analysis of protein oligomerization and hydrophobicity

Oligomerization state of AH3-GFP is first analyzed by protein concentration tube Vivaspin 2 with MWCO of 100, 300, or 1000 kDa from Sartorius. Protein samples were centrifuged in Vivaspin 2 at 1000g for 20 minutes. Filtrate was analyzed by SDS-PAGE and then coomassie blue staining for analysis. TEM images of AH3-GFP protein complex was obtained after negative staining with phosphotungstic acid.

Images were taken using Tecnai G2 Spirit Twin. For AH3-sfGFP-2xhM2e and its variants, the purified fusion protein was first crosslinked with Sulfo-SMCC(sulfosuccinimidyl 4-[N-maleimidomethyl]cyclohexane-1-carboxylate) at 4 μg/ml for 30 minutes before been processed using negative staining protocol. A sucrose density gradient was used to analyze AH3-sfGFP-2xhM2e related proteins oligomerization status and hydrophobicity. In a 13-ml polypropylene tube (Beckman cat#14287), the bottom was layered first with 1 ml 65% (w/v) sucrose solution and then was topped with 2 ml 45% (w/v) sucrose solution and then followed by 7 ml 15% (w/v) sucrose solution. Sudan III stock solution was prepared as 0.5% in isopropanol. Staining of bacterial membrane was by adding Sudan III stock solution into bacterial suspension before sonication at 1:100 ratio. One milliliter of the soluble fraction was layered on top of 15% sucrose solution. To test protein nanoparticle membrane binding activity, 1.5ml baterial lysate from empty vector transformed BL21(DE3) was mixed with 0.3mg purified protein nanoparticle and incubated in RT for 30 minutes before been layered on the top of sucrose solution and ultracentrifugation. The centrifuge tubes were photographed in front of dark background by exposed to 450 nm LED light. Images were processed in ImageJ by first extracting green channels, then the fluorescent intensity of ultracentrifugation results were quantitated from top to bottom using Plot Profile and compiled using ROI manager. Protein nanoparticle hydrodynamic diameter is determined by dynamic light scattering (DLS) using Stunner (Unchained Lab).

### Animal immunization and antibody titer determination

The animal protocols had been approved by IACUC of Fu Jen Catholic University and mice were housed in the experimental animal center of Fu Jen Catholic University following SPF standards. Mice used in immunization procedures were between 8 to 9 weeks age. Mice were immunized through an intramuscular injection of purified protein preparation in 1 mg/ml concentration. For blood collection, mice were bled from a facial vein after pricked by the lancet. Sera collected were stored in −80 °C before been analyzed by ELISA assay. For each immunization, 20 μg purified protein was injected intramuscularly in the thigh of the hind limb. Blood was bled 14 days post immunization or planned dates for analysis using ELISA. For ELISA, the antigen was diluted with coating buffer for coating on high binding ELISA plate (Greiner Bio-One MICROLON high binding ELISA plate), antigen concentration of hM2e peptide was 2.5μg/ml and sfGFP was 1 μg/ml. The ELISA plate was incubated at 4 °C overnight and then washed and blocked with 200 μl of blocking buffer containing 1% BSA in washing buffer. Washing buffer is composed by 10mM Na(PO_4_), 150mM NaCl at pH 7.4 with 0.05% Tween 20. Antiserum was diluted started from 1:100 and followed by 4 folds serial dilution. Secondary antibody, Goat anti-mouse IgG with HRP conjugation was diluted at 1:5000 in blocking buffer.

Chromogenic development was carried out by adding 100μl TMB and incubated for 10 minutes and stopped by adding 100μl 2M H_2_SO_4_. Antibody titer was determined as the reciprocal value of the highest dilution that gives an OD450 reading of 0.1 above background.

### Protein structure modeling and intermolecular force calculation

Protein structure models of AH3 peptide monomer, dimer and tetramer were generated using AscalaphDesigner version 1.8.79 and manual movement. The intermolecular interaction forces between either monomers or dimers were calculated using the intermolecular energy command of AscalaphDesigner. Hydrogen bonds between peptide subunit and hydrophobic patch were determined and illustrated using Deep View/Swiss PDB Viewer version 4.1.1. The solvent accessible surface area of the AH3 dimer and GFP were calculated using Jmol with a radius of 1.2 angstroms.

#### Thermostability determination

The purified protein nanoparticle stored in elution buffer was changed to 0.5x GF buffer pH 8.0 using Sephadex-25 resin. Then the NaCl concentration of protein solutions was adjusted to designed concentration by adding 5M NaCl. Protein samples were incubated in various temperatures and samples were taken at certain time points for sample preparation for SDS-PAGE and coomassie blue staining at the end.

## Results

### Identification of a protein complex with high antigenicity and stability

As described in our patent application filed in 2015, we have tested the immunogenicity of fusion proteins composed of an AHP and a GFP (Kan, 2018). The results showed an increase of anti-GFP IgG titer ranged between 2∼3 log under a two immunizations regime (Figure S1). One of the peptides, AH3, that derived from M2 protein of type A influenza strain H5N1 gives extended stability to the GFP fusion protein when compared to another peptide, AH1 (Figure S2) as well as other peptides in our study (data not shown). Since a stable protein is essential for vaccine carrier, we were interested in the mechanism of AH3-GFP stability and antigenicity. To study the potential mechanisms that contribute to the above mentioned properties of AH3-GFP fusion protein, we first checked the composition of AH3-GFP protein post expression and purification. One clue that led us to study the composition of AH3-GFP fusion protein is the difficulties encountered during protein purification. Unlike other fusion proteins studied, most of the AH3-GFP and AH5-GFP fusion proteins are expressed as insoluble inclusion body and the remaining soluble protein did not bind to Ni-NTA resin under normal condition of 300mM NaCl. The AH3-GFP and AH5-GFP fusion proteins only started to bind to Ni-NTA resin after we decreased the NaCl concentration from 300mM to 50mM. Also, the resistance of AH3-GFP fusion protein to hydrolysis suggested the linker between AH3 peptide and GFP is kept in a water tight complex. The likely hypothesis to explain these observations is when GFP was fused with AH3 peptide, the AH3 peptide induces the assembly of a hydrophobicity driven protein complex that hinders N-terminal His tag from binding to Ni-NTA ligand.

### Characterization of AH3-GFP protein complex

To test the hypothesis that AH3-GFP or AH5-GFP fusion protein forms a protein complex, we first used protein concentration tube with different molecular weight cut off (MWCO) to determine the protein complex sizes. As shown in figure 1A, GFP protein with a molecular weight of 27kDa was able to pass through membranes with MWCO of 100kDa, 300kDa and 1000kDa freely, but AH3-GFP fusion protein was prevented from passing through the membrane with an MWCO up to 1000kDa. With a molecular weight of 30kDa, the purified AH3-GFP fusion protein need to form a complex with more than 35 monomers to be excluded from passing a membrane with a 1000kDa MWCO. To explore further the geometric composition of the AH3-GFP protein complex, we examined the fusion protein under transmission electronic microscope (TEM). The TEM imagess showed the AH3-GFP fusion protein forms a corn-on-a-cob structure with a length up to 60 nm and a diameter around 10 nm (Fig. 1B). The difference in length suggests that the particle may be assembled along the long axis. When scanning along the long axis of the AH3-GFP particle, there is a repetitive pattern of two-one-two-one of white dots with less visible dot on each side of the single dot. The predicted structure according to TEM images is shown in Fig.1D. We also examined AH5-GFP protein complex under TEM, but there is no clear evidence of forming higher order protein complex, suggesting AH5-GFP protein complex is not as stable as AH3-GFP to withstand the conditions during negative staining. To find the correlation between protein complex formation and antigenicity, we immunized mice with purified AH3-GFP fusion protein and the recombinant GFP protein. Proteins were prepared from LPS synthesis defective E. coli strain, ClearColi BL21(DE3), to avoid the interference of LPS contamination, a known TLR4 ligand and adjuvant. The mice were immunized with purified proteins by single intramuscular injection and sera were collected at day 7, 14, 30 and 182 to evaluate anti-GFP IgG titer by ELISA. Deoxycholate was added to test if deoxycholate in the concentration of 0.2% affects AH3-GFP antigenicity and related experiment was terminated at 30 days post immunization when it showed no effect on antigenicity of either GFP or AH3-GFP. These results suggest GFP alone is a poor antigen and only gained high antigenicity after fused with AH3 peptide and forms protein nanoparticle (Fig. 1C).

**Figure 1.**
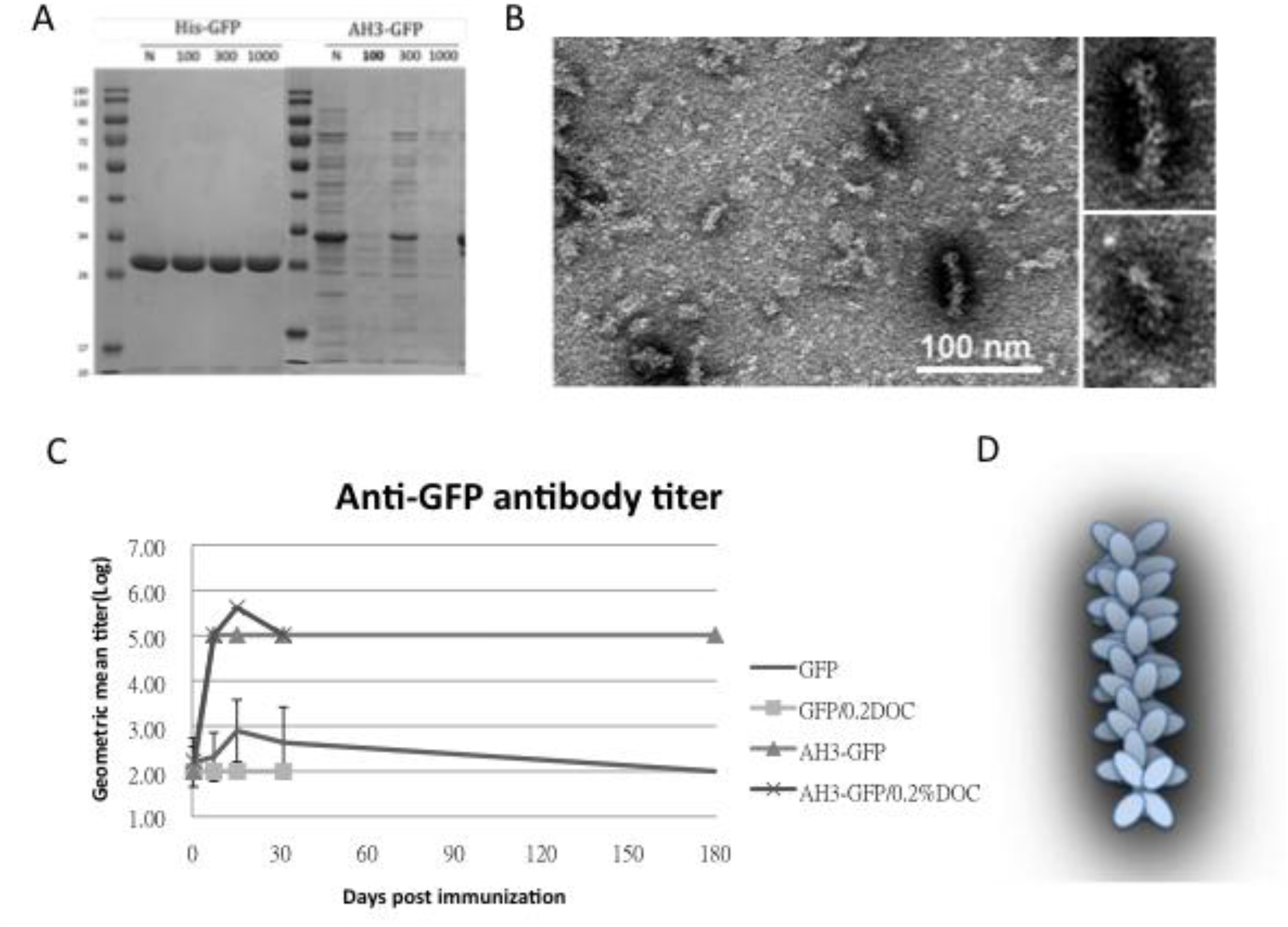
Characterization of GFP fusion protein oligomerization. (A) GFP and AH3-GFP fusion proteins purified and reconstituted in PBS were centrifuged through size exclusion membrane with different molecular weight cut off (MWCO). The filtrate was then analyzed by 12% SDS-PAGE and coomassie blue staining (arrows are needed to point out the bands). (B) Antigenicity of GFP and AH3-GFP fusion protein were evaluated by immunizing mice by single muscular immunization. The anti-GFP IgG titers were followed for 6 months by ELISA. (N=5). Two samples (GFP/0.2% DOC and AH3-GFP/0.2%DOC) also included 0.2% deoxycholate when immunizing mice. (C) The oligomerization of AH3-GFP protein was analyzed by negative staining using a transmissive electronic microscope(TEM). Scale bar represents 100nm. Two of the representative particles are enlarged and shown in the right panels. (D) AH3-GFP protein nanoparticle model predicted from TEM images.

### Modeling AH3-GFP protein complex structure

To understand the potential molecular mechanism leading to the assembly of AH3-GFP nanoparticle, we built an AH3 models based on the known helical structure and the hypothesis that hydrophobic interaction drives the complex formation in addition to two observations from TEM images: first, the particle was assembled along the long axis and the second, AH3-GFP protein particle has a repetitive three-two pattern along long axis after examining protein particle images. We first assembled two AH3 peptides as anti-parallel helices with a 24 degree angle to make close contacts involving side chains of Phe5, Phe6, Ile9, Leu13 and Leu17 using protein modeling software, AscalaphDesigner (Fig. 2A). The hydrophobic core contributes the major intermolecular force binding two helices and the AH3 dimer is surrounded by hydrophilic side chains from multiple lysine and arginine. Using another protein modeling software, Deepview, a hydrophobic patch was identified on one face of the assembled dimer as marked by red surface (Fig. 2B). In a water accessible surface model of AH3 dimer, two connected hydrophobic pockets can be seen located within the hydrophobic patch (marked by dash line) that serves as binding sites for two Arg12 side chains extruding from the opposite face of second AH3 dimer. The second AH3 dimer can make close contact in tandem with the first dimer after turning counter clockwise for 36° looking down the hydrophobic patch and forms a tetramer (Fig. 2C). The intermolecular energy between two dimers from this model was calculated to have a ΔG of −101 kcal/mol (Fig. 2C). After adding GFP protein structures onto the AH3 tetramer model, the AH3-GFP fusion protein tetramer forms a scissor-shaped assembling unit and the stacking of every AH3-GFP tetramer on top of another tetramer will increase the particle length by 2.8 nm and turning the axis by 72°. Since the GFP protein barrel diameter is ranged between 2.7∼3.5 nm, the out extending GFP from AH3-GFP tetramer can spatially fit into this model (Fig. 2D). The length of linker between AH3 peptide and sfGFP is important for protein nanoparticle assembly as well since fusion protein with a 2 amino acids linker failed to assemble into nanoparticle. According to this model, the protein nanoparticle will be extended continuously along the long axis with a hydrophobic patch presenting on the growing end of the assembling particle and serving as a point for polymerization.

**Figure 2.**
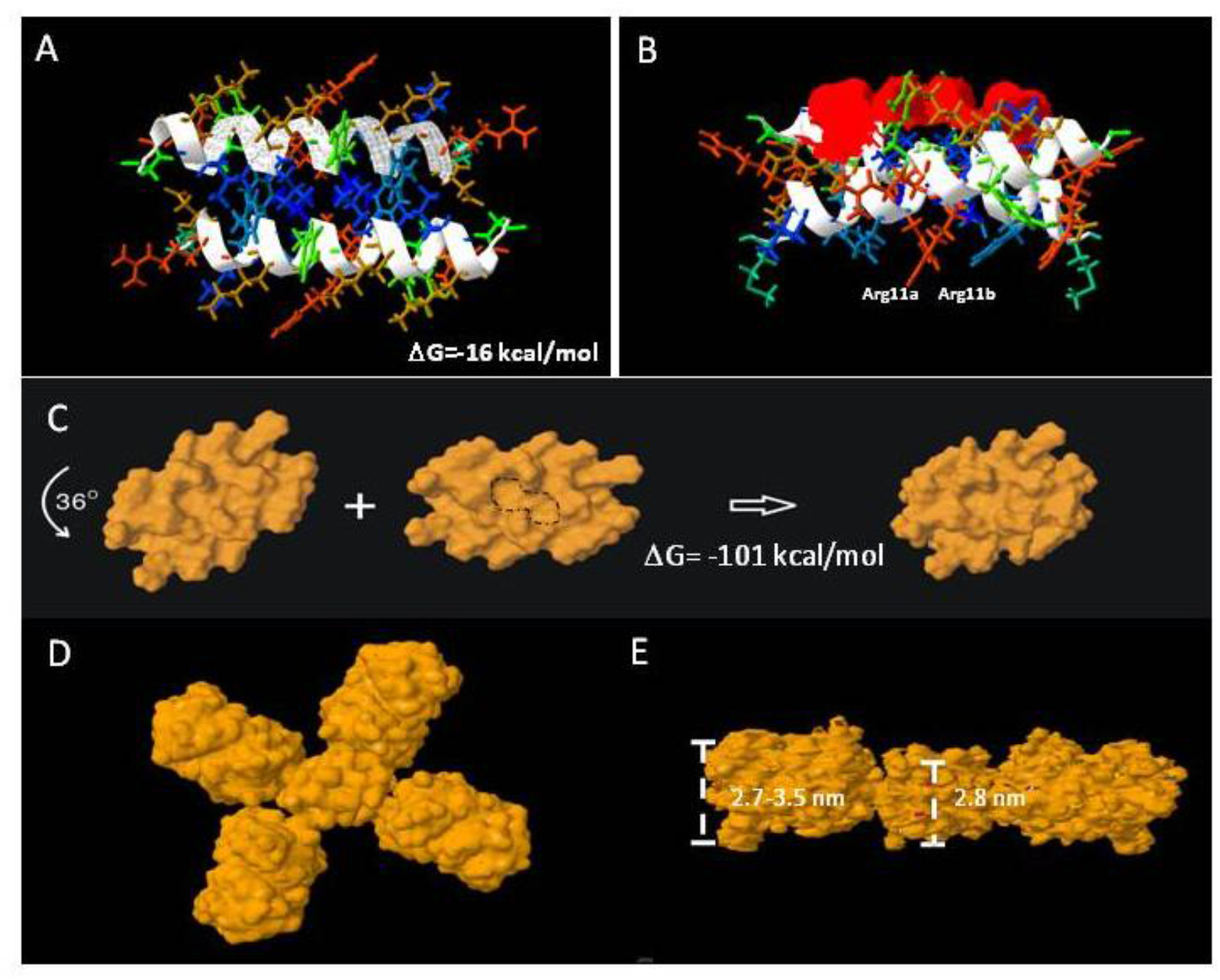
AH3-GFP protein nanoparticle structural modeling. (A) The assembling unit of AH3-GFP protein nanoparticle was modeled as dimer interacting through hydrophobic interaction using AscalaphDesigner. Amino acid side chains are colored according to hydrophobicity using SPDB v4.10. The most hydrophilic side chains are colored red and most hydrophobic side chains are colored blue, side chains with hydrophobicity in between are colored transitionally. The two α-helix backbones are marked by white ribbon. The intermolecular interaction force (ΔG) is shown in the bottom right corner. (B) Bottom view of AH3 dimer that shows the location of the hydrophobic patch (marked by red surface). The anti-parallel alpha helix from two monomers forms a cross and has an angle of 24 degrees. (C) Water accessible surface of AH3 dimer is modeled using Pymol and the stacking of 2^nd^ dimer onto the first one is achieved after turning the 2^nd^ dimer counter clockwise for 36°. The two hydrophobic pockets are enclosed in dash line. The intermolecular force (ΔG) between two dimers is shown. (D)The AH3 tetramer with 4 fused GFP molecules is modeled to form a cross shape. (E) The bottom view of AH3-GFP tetramer is shown with two GFP molecules removed for a clear view. The distance between two repeating atoms of stacked tetramer is measured and shown as the thickness of AH3 tetramer, the thickness of GFP monomer is measured as the distance across the protein barrel structure.

### Designing a vaccine carrier that enables heterologous antigen insertion and high stability

After proving that the AH3-GFP protein nanoparticle possesses high antigenicity, we decided to explore the potential of AH3-GFP protein nanoparticle as a vaccine carrier. For the Hepatitis B core antigen, the amino acid 144 may serves as an insertion site for heterologous antigen fusion and exposes the antigen to immune system (Borisova et al., 1989). GFP protein has a thermal stable structure that constituted by 11 β-strands and 1 α-helix and some of the loops connecting β-strands have been explored as insertion sites for heterologous protein(Kobayashi et al., 2008; Pavoor et al., 2009; Wang et al., 2014). Among those candidates, loop173 linking strand 8 and strand 9 was chosen because it had been shown to has a high capacity for foreign peptide insertion (Fig 3A) (Kobayashi et al., 2008). The original AH3-GFP recombinant protein is constructed in pET28a vector with AH3 coding sequence inserted C-terminal to His-tag and thrombin cleavage site followed immediately by GFP cloned from pEGFP-C2. This expression vector was low in soluble protein production and unable to express soluble recombinant protein when the peptide is inserted between D173 and G174. To resolve the expression and folding issues, we designed a new expression vector. First, we cloned AH3 peptide into the very N-terminal following methionine in the pET27 vector, and then to its C-terminal we inserted a 6 a.a. linker (GTTSDV) followed by a synthetic sfGFP gene (Pédelacq et al., 2006) with an antigen insertion site next to Ser175 of sfGFP. The antigen insertion site also contains an 8xHis tag to facilitate protein purification. To verify vaccine carrier function, we inserted two copies of a broad spectrum flu vaccine candidate, M2 ectopic peptide from PR8 strain (hM2e) separated by a 6 a.a. linker (Fig. 3B). The newly constructed vector was proven to be efficient for expressing soluble AH3-sfGFP-2hM2e fusion protein as a protein complex. But the AH3-sfGFP-2xhM2e protein complex is not as stable as AH3-GFP because these nanoparticles fall apart during distilled water wash before TEM imaging, after few trials we found the protein nanoparticles have to be crosslinked by sulfo-SMCC before TEM analysis (Fig. 5A). Following the established AH3-GFP protein complex model, we were seeking strategies to create a more stable AH3-sfGFP protein complex. First, using protein modeling software AscalaphDesigner, we found the mutation of Ile9 to Leu increases the intermolecular force (ΔG) between two peptide helices from −16 kcal/mol to −39 kcal/mol (Fig.3C). Second, when we mutated Lys14 to Glu14, there are two hydrogen bonds forms between LYRRLE dimers, one between the side chain of Glu14c and Cys8a and the other one forms between side chain of Arg12a and the oxygen of Tyr10c backbone (Fig. 3D). The intermolecular force between the LYRRLE dimers increase from −101 kcal/mol to −161 kcal/mol. The K14E mutation not only enables tighter binding between two adjacent dimers, but also resolves a key issue of AH3 mediated protein nanoparticle assembly, an exposed hydrophobic patch. From the protein modeling results, we found that the dimer assembled from AH3 double mutant, LYRRLE, is able to bind to preformed particle in two orientations: either as a tandem dimer (ΔG=-161 kcal/mol)(Fig. 3F) as an inverted dimer (ΔG=-59 kcal/mol). The formation of inverted tetramer is able to enclose the hydrophobic patch inside the protein nanoparticle (Fig. 3G).

**Figure 3.**
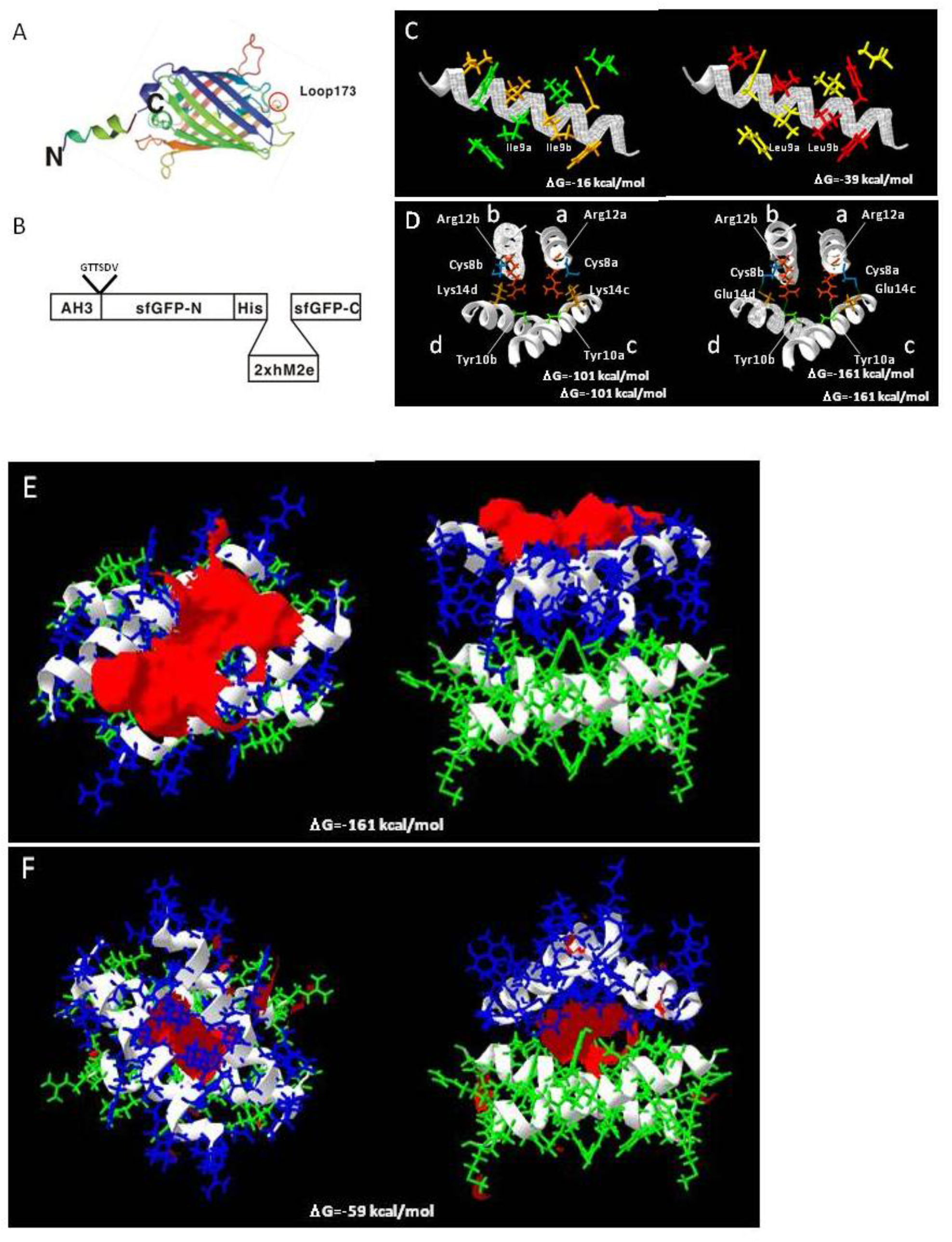
Construction of AH3-sfGFP-2xhM2e fusion protein and protein model guided AH3 mutagenesis for variants with a higher nanoparticle stability. Graphic presentation of AH3-sfGFP fusion protein is shown and the antigen insertion site (loop173) is marked by a red circle. The 19 amino acid AH3 peptide was fused to the N-terminal of sfGFP to mediate polymerization and a foreign antigen insertion site is genetically created in loop173 containing an 8xHis tag for efficient protein purification. (B) The graphic presentation of AH3-sfGFP-2xhM2e fusion protein is shown. The antigen insertion site can accommodate antigenic peptide with size up to 100 a.a. (C) Comparison of AH3 and AH3 I9L (LY) mutation in dimer formation. The intermolecular force between two monomers is shown. (D) The hydrogen bonds form between AH3 variant LYRRLE dimers are shown in the right panel. AH3 tandem tetramer is shown in the left panel. Four monomers from each model are labeled from a-d. The Glu14 from monomers c and d form hydrogen bonds with Cys8 from monomers a and b respectively. Also the side chain of Arg12 from a and b monomers forms hydrogen bond with back bone of monomer c and d respectively. (E) The front view (left panel) and bottom view (right panel) of LYRRLE tandem tetramer protein models. (F) The front view (left panel) and bottom view (right panel) of LYRRLE inverted tetramer. The hydrophobic patch is marked as red surface, the first dimer is colored as green and second dimer is colored as blue. The calculated intermolecular interaction force is shown in the center of both graphs.

Following the protein modeling result, we go further to verify whether the protein modeling results are correct using biophysical assay. We generated mutations in AH3 peptide sequence in the context of AH3-sfGFP-2xhM2e (Fig. 4A). Our hypothesis is that the exposed hydrophobic patch on the end of the AH3-GFP protein nanoparticle will bind bacterial membrane and co-sediment with membrane during sucrose step gradient ultracentrifugation. This methodology was first verified by loading bacterial soluble fraction (containing bacterial membrane) prepared from sonicated ClearColi bacterial culture in a centrifuge tube preloaded with 15%(w/v), 45%(w/v) and 65%(w/v) sucrose as demonstrated in Fig. 4B left panel. The sedimentation of bacterial membrane was marked by a lysochromic dye (Sudan III) that binds lipid.

**Figure 4.**
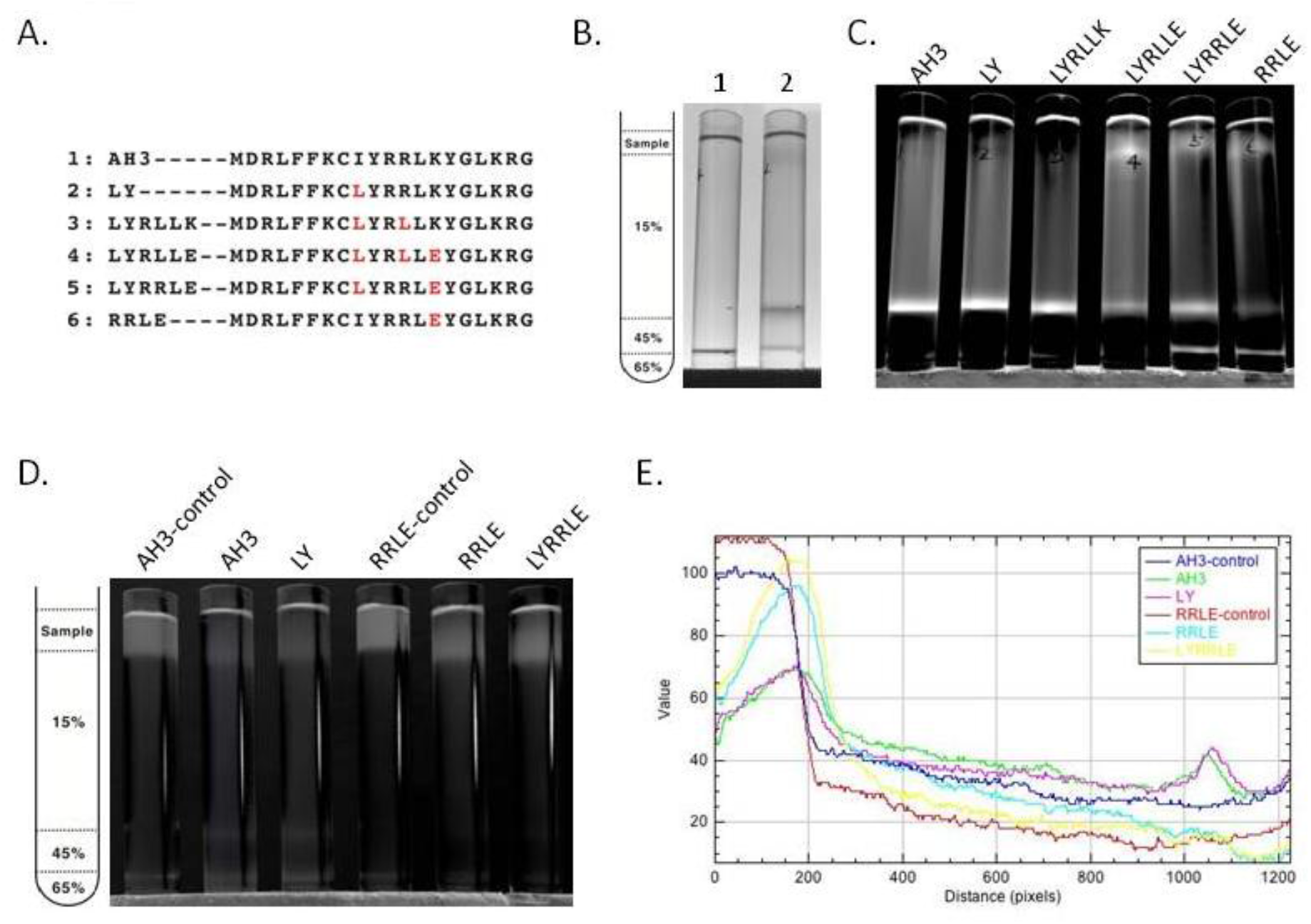
The co-sedimentation analysis of AH3-sfGFP-2xhM2e variants with bacterial membrane by analytic ultracentrifugation. (A) List of AH3 peptide variants introduced on AH3-sfGFP-2xhM2e fusion protein for co-sedimentation analysis. (B) The right panel shows the graphic presentation of the centrifuge tube distribution of step sucrose solutions. The left panel shows the bacterial membrane distribution after ultracentrifugation as marked by Sudan III staining. Lane 1 is a Sudan III solution only control without bacterial lysate, lane 2 is topped with 1ml bacterial lysate mixed with Sudan III solution. The ratio between Sudan III staining solution and bacterial lysate is 1:100. (C) The distribution of sfGFP fusion proteins post ultracentrifugation as illuminated by 450nm LED light was recorded as photo and analyzed by ImageJ. Only the green channel is shown. (D) The ability of purified protein nanoparticle to bind bacterial membrane were analyzed by first mixing with bacterial lysate and then analyzed by ultracentrifugation on step sucrose gradient. The distribution of sucrose solutions of different percentages is shown in the left panel. The protein nanoparticle purified from fusion proteins containing various AH3 mutants, AH3, LY, RRLE and LYRRLE were tested. The control samples did not go through ultracentrifugation to show the original position of loaded samples. The photo taken is split into three channels by ImageJ, only green channel is shown. (E) The combined line plot data of fluorescent intensity along the centrifuge tube from top to bottom. The image pixels had been averaged by the “smooth” command of imageJ before line plot analysis to decrease variations.

The control sample contained Sudan III with lysis buffer alone (Fig. 4B lane 1). After ultracentrifugation, the bacterial membrane was sedimented to the junction between 15% and 45% sucrose as marked by Sudan III staining(Fig.4B lane 2). Using the same protocol, AH3-GFP was found to co-sedimented with the bacterial membrane (Fig. S3A) as well as the AH3-sfGFP-hM2e fusion protein but not a free GFP protein (Fig S3B) when the tubes were illuminated by 450nm LED light. After protocol verification, the bacterial soluble fraction from all six fusion protein expression cultures were prepared and analyzed using the same protocol. From the distribution of fluorescent protein between different percentage of sucrose, we have made several observations: first, all of expressed AH3-sfGFP-2xhM2e protein binds to bacterial membrane and co-sedimented to 15%/45% junction, so as AH3 variants LY and LYRLLK (Fig. 4C lane 1). Second, comparing tubes 4, 5 and 6 to tubes 1, 2 and 3, it is clear that when Lys13 is mutated to Glu, this mutation decreases the binding of expressed protein nanoparticles to bacterial membrane and about half of the protein complex remained on the top of centrifuge tube. Third, the protein complex of AH3 variant LYRRLE and RRLE even forms higher order protein aggregate and was sedimented further to 45%/65% junction (Fig. 4C lane 5 and 6), the identity of these high order protein complex is not known, most likely it was aggregated protein nanoparticles. These results confirmed the predicted presence of a hydrophobic patch on protein nanoparticle assembled from AH3-sfGFP-2xhM2e protein and LY, LYRLLK variants. And a portion of the protein nanoparticle assembled from AH3 variants containing a K13E mutation removed the hydrophobic patch from surface when they were synthesized in cell. To verify if purified protein nanoparticle still poses membrane binding activity, the purified protein nanoparticle of AH3, LY, RRLE and LYRRLE variants were first mixed with bacterial soluble fraction and incubated for 30 minutes before been evaluated by ultracentrifugation. A control tube with same setup but no centrifugation was used to show the position of protein nanoparticle before ultracentrifugation (Fig. 4D). The data shows, a fraction of protein nanoparticle from AH3 and LY variants bind and co-sediment with bacterial membrane during ultracentrifugation as shown in line plot that depicts fluorescent protein distribution of all test samples (Fig. 4E). None of the protein nanoparticle assembled by RRLE or LYRRLE variants co-sedimented with bacterial membrane. These data is consistent with the protein modeling results that protein nanoparticle assembled by LYRRLE variant is able to eliminate hydrophobic patch through an inverted cap (Fig. 3F).

To further evaluate the effect of AH3 mutants on protein nanoparticle formation, we examined the particle morphology using TEM and Dynamic light scattering (DLS). The result is shown in the negative staining images (Fig. 5A-D). Through visual observation and image refinement using ImageJ, we found some morphological evidence that support protein modeling result. First, there is donut-shape protein nanoparticle been observed in AH3-sfGFP-2xhM2e protein prep (Fig. 5A). The donut or disc shape structure may be due to collapse of corn-on-a-cob structure because of low intermolecular interaction between AH3 monomers (ΔG= −16kcal/mol). This hypothesis is supported by the DLS data that measures the protein particle size. These data show a 2.5 folds increase in particle size when Ile8 (AH3) is mutated to Leu8 (LY) and an increase of predicted intermolecular force between monomers from ΔG= −16kcal/mol to ΔG= −39kcal/mol (Table 1). The significant of Leu8 in stabilizing particle structure is also underscored by the change in morphology of protein nanoparticles assembled by RRLD and RRLE variants. Part of the fusion proteins containing RRLD or RRLE variants assembled into eccentric ladder-like structure, suggesting an imbalance in the interaction forces within protein nanoparticle.

**Figure 5.**
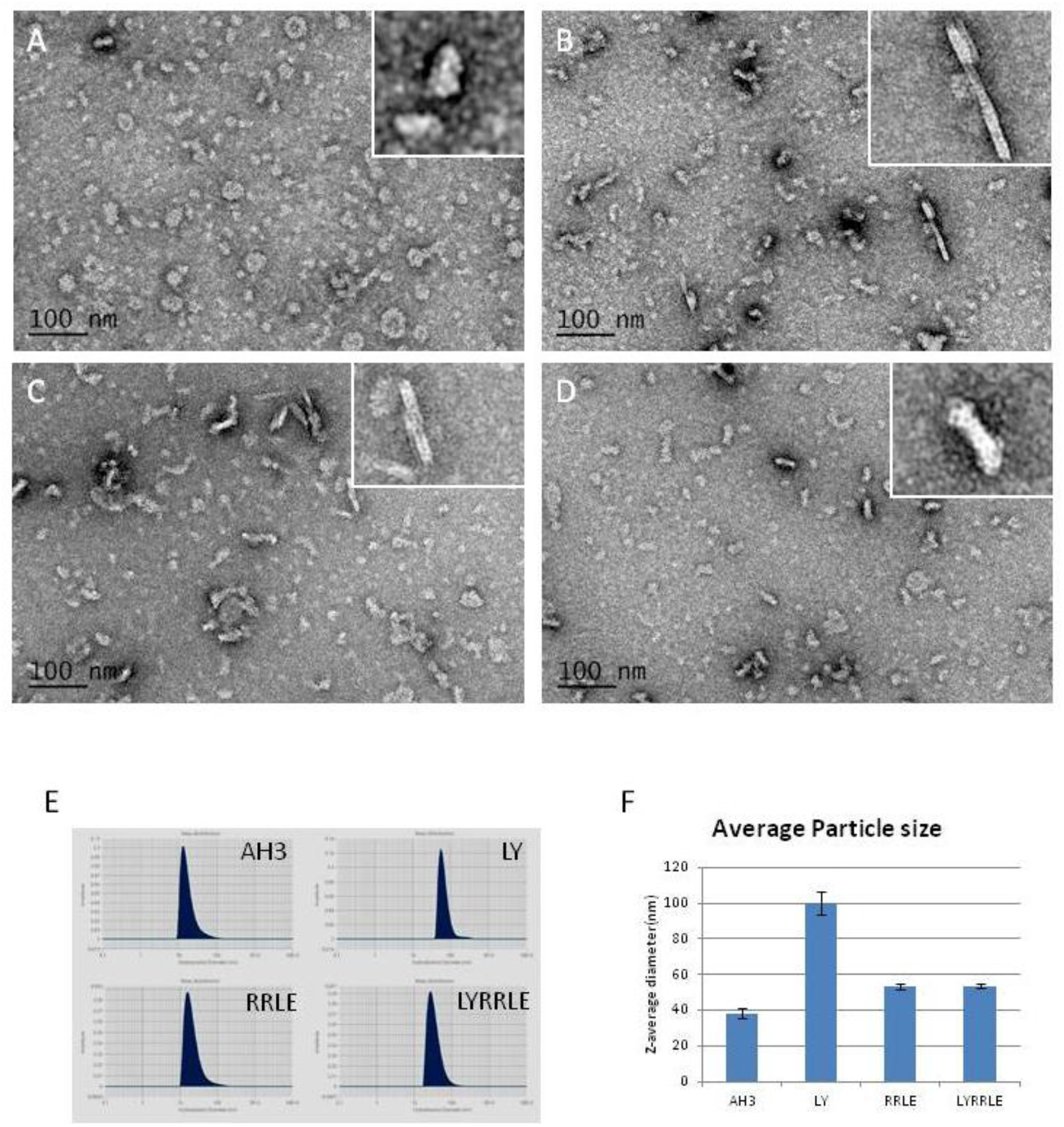
The TEM images of protein nanoparticles derived from AH3-sfGFP-2xhM2e and its variants. Purified protein nanoparticles were crosslinked with Sulfo-SMCC after buffer change from elution buffer using desalting resin (Sephadex 25). Protein samples were diluted to 0.1mg/ml before applied on the grid for TEM analysis. The representative images were presented in the sequence of (A) AH3-sfGFP-2xhM2e, (B) RRLD-sfGFP-2xhM2e, (C) RRLE-sfGFP-2xhM2E AND (D) LYRRLE-sfGFP-2xhM2e. The enclosed window on the upper-right corner shows the magnified images of representative protein nanoparticle. The scale bar of 100nm is included. (E) The hydrodynamic diameter of AH3-sfGFP-2xhM2e derived protein nanoparticle was analyzed by dynamic light scattering (DLS). (F) The Z average diameter of AH3-sfGFP-2xhM2e derived protein nanoparticles are compared.

**Table 1.**

|  | Assembling unit | Intermolecular interactions(kcal/mol) | Hydrophobic patch area (Å<sup>2</sup>) |
| --- | --- | --- | --- |
| AH3 | Monomer-Monomer | -16 | 285 |
|  | Tandem dimer-dimer | -101 | 386 |
|  | Inverted dimer-dimer | na | na |
| LY | Monomer-Monomer | -39 | 389 |
|  | Tandem dimer-dimer | na | na |
|  | Inverted dimer-dimer | na | na |
| RRLE | Monomer-Monomer | -23 | 429 |
|  | Tandem dimer-dimer | -29 | 342 |
|  | Inverted dimer-dimer | na | na |
| LYRRLE | Monomer-Monomer | -36 | 393 |
|  | Tandem dimer-dimer | -161 | 433 |
|  | Inverted dimer-dimer | -59 | 111 |

As shown in the sucrose step gradient ultracentrifugation, the presence of hydrophobic patch enables nonspecific interaction of the AH3-GFP protein complex with bacterial membrane. Which may restrict the free moving of protein nanoparticle and keep it from reaching draining lymph node for stimulating immunity (Bachmann and Jennings, 2010). To compare the antigenicity of protein complexes derived from either AH3-sfGFP-2xhM2e or LYRRLE-sfGFP-2xhM2e, we immunized mice with a single injection of either recombinant proteins. Post immunization, sera were collected at day 15, 50, 90 and 202 to evaluate anti-hM2e IgG titer by ELISA. The geometric mean titer of anti-hM2e IgG reached the highest point for the AH3-sfGFP-2xhM2e group and then declined afterward. But of the LYRRLE-sfGFP-2xhM2e group, the GMT reached highest point at day 50 and remained steady up to day 90 (Fig. S4). When the individual mouse serum result is observed separately, only one out of 5 mice from AH3-sfGFP-2xhM2e group has higher anti-hM2e IgG titer at day 202 than day 15. But there are 4 out of 5 mice from the LYRRLE-sfGFP-2xhM2e group shows a higher antibody titer in day 202 compared to day 15 (Fig. 6A). These results suggest that the two point mutations of AH3 in I9L and K14E enable the formation of a stable, high antigenic protein complex that stimulates long lasting immune responses in a single immunization.

**Figure 6.**
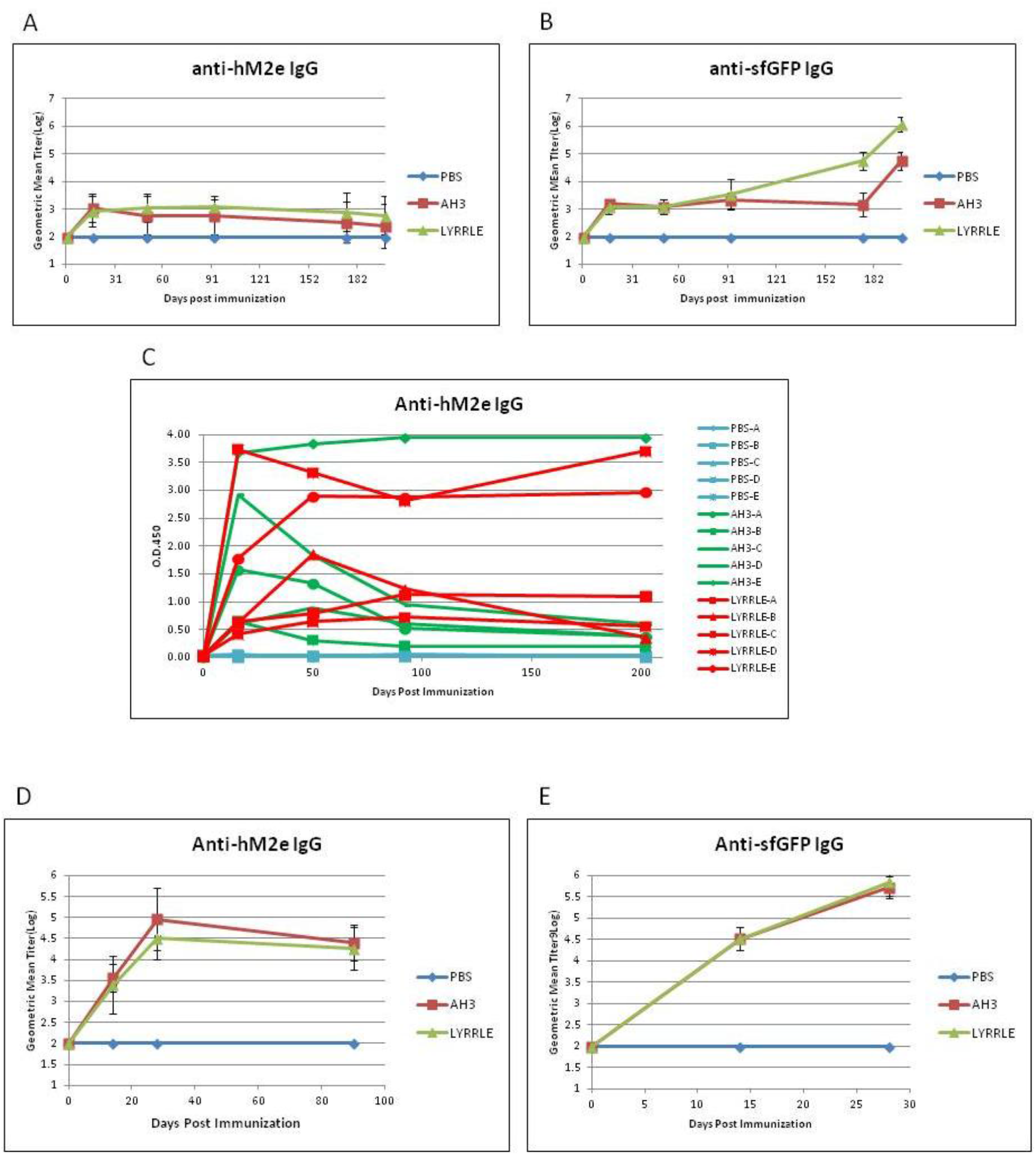
Immunization of mice with AH3-sfGFP-2xhM2e and LYRRLE-sfGFP-2xhM2e and detection of anti-hM2e IgG and anti-sfGFP IgG. The mice were immunized once at day 0 by 20 μg of purified proteins nanoparticle assembled by wild type AH3 peptide or LYRRLE mutant or PBS. Sera were collected at day 0, 16, 50, 991, 175 and 199 for analysis to detect (A) total anti-hM2e IgG and total anti-sfGFP IgG by ELISA. (C) Anti-hM2e total IgG titers are presented as optical density at OD450 using immune serum from each mouse diluted 1:100. (D) The anti-hM2e total IgG of mice immunized twice with 14 days apart in a prime-boost regime were followed for 3 months and analyzed by ELISA (N=5). (E) The anti-sfGFP total IgG of mice immunized by AH3-sfGFP-2xhM2e or LYRRLE-sfGFP-2xhM2e twice were evaluated 14 days post each immunization by ELISA (N=5).

Vaccine carrier like HBc based virus like particle (VLP) is often failed at boosting humoral immune responses after prime dose in a multiple doses protocol due to antigen competition. Although GFP is a protein of low antigenicity, the fusion with AH3 strongly enhances its antigenicity as shown in Figure 1C. To test if sfGFP backbone competes with inserted hM2e peptide for immune machinery, we immunized mice in a prime-boost protocol using the same protein preparations. The two consecutive injections were carried out 14 days apart and sera collected at day 14, 28 and 90 were subject to ELISA assay using either hM2e peptide or sfGFP protein as coating antigen. The result shows the IgG titer against hM2e elevated continuously after consecutive immunizations for both proteins as well as anti-sfGFP IgG titer. The result suggests that although carrier protein AH3-sfGFP also has high antigenicity, it does not interfere with the immune response against the heterologous protein, hM2e (Fig. 6B, 6C).

One possible explanation of long lasting antibody response is the continuous stimulation of immune system by the protein nanoparticle assembled by LYRRLE peptide but not AH3 peptide. To test this hypothesis, we incubated the purified protein nanoparticle in phosphate buffer containing 150 mM NaCl at 40°C or 50 °C up to 4 weeks. The integrity of protein nanoparticle was examined by SDS-PAGE. The data suggest protein nanoparticle assembled by LYRRLE and RRLE have better thermostability than AH3 and LY peptides. This result is consistent with the protein modeling result that shows a hydrogen bond forms between two dimers when Lys13 was mutated to glutamic acid (Fig. 3D). Combining the above data, we find the LYRRLE peptide is more suitable in serving as a vaccine carrier because two reasons, first: it is more stable in host body temperature so it can endure high temperature storage; second: the removal of hydrophobic patch from surface enables the nanoparticle move freely through vessels to reach target site.

During manipulation of LYRRLE protein nanoparticle, it was observed that protein nanoparticle often degraded within short storage time after buffer was changed from high salt elution buffer (300mM NaCl, 500mM Imidazole) to isotonic buffer (150mM NaCl). This suggests a possibility of salt concentration dependent nanoparticle stabilization. To evaluate the effect of NaCl concentration on the stability of LYRRLE assembled protein nanoparticle, the LYRRLE-sfGFP-2xhM2e protein nanoparticle was incubated in 4 °C, 22 °C and 37 °C up to 4 weeks and protein stability was evaluated by SDS-PAGE.

When protein nanoparticle was stored in 4 °C, the protein in buffer containing 150mM NaCl was stable at first week, but hydrolyzed into lower molecular weight before 4^th^ week (Fig. 7B). And protein nanoparticle in buffer containing 300mM NaCl was stable for one month but it was later hydrolyzed before the end of third month (Fig. 7C). When the storage temperature is raised to 22 °C, the protein was hydrolyzed within a week in buffer containing 150mM NaCl and within 4 weeks in buffer containing 150, 300 and 550mM NaCl. The protein nanoparticle is only stable in buffer containing 800mM NaCl (Fig. 7D). When the temperature was shifted to 37 °C, the 300 mM NaCl is sufficient to keep protein nanoparticle intact in the end of 3 months incubation (Fig. 7E, 7F). To be noted is the protein samples that contain hydrolyzed protein nanoparticle were still fluorescent, an indication that sfGFP is still functionally intact. According to the size of hydrolysis product (20kDa), the protein hydrolysis is likely happened to the inserted loop that contains 2xhM2e peptide.

**Figure 7.**
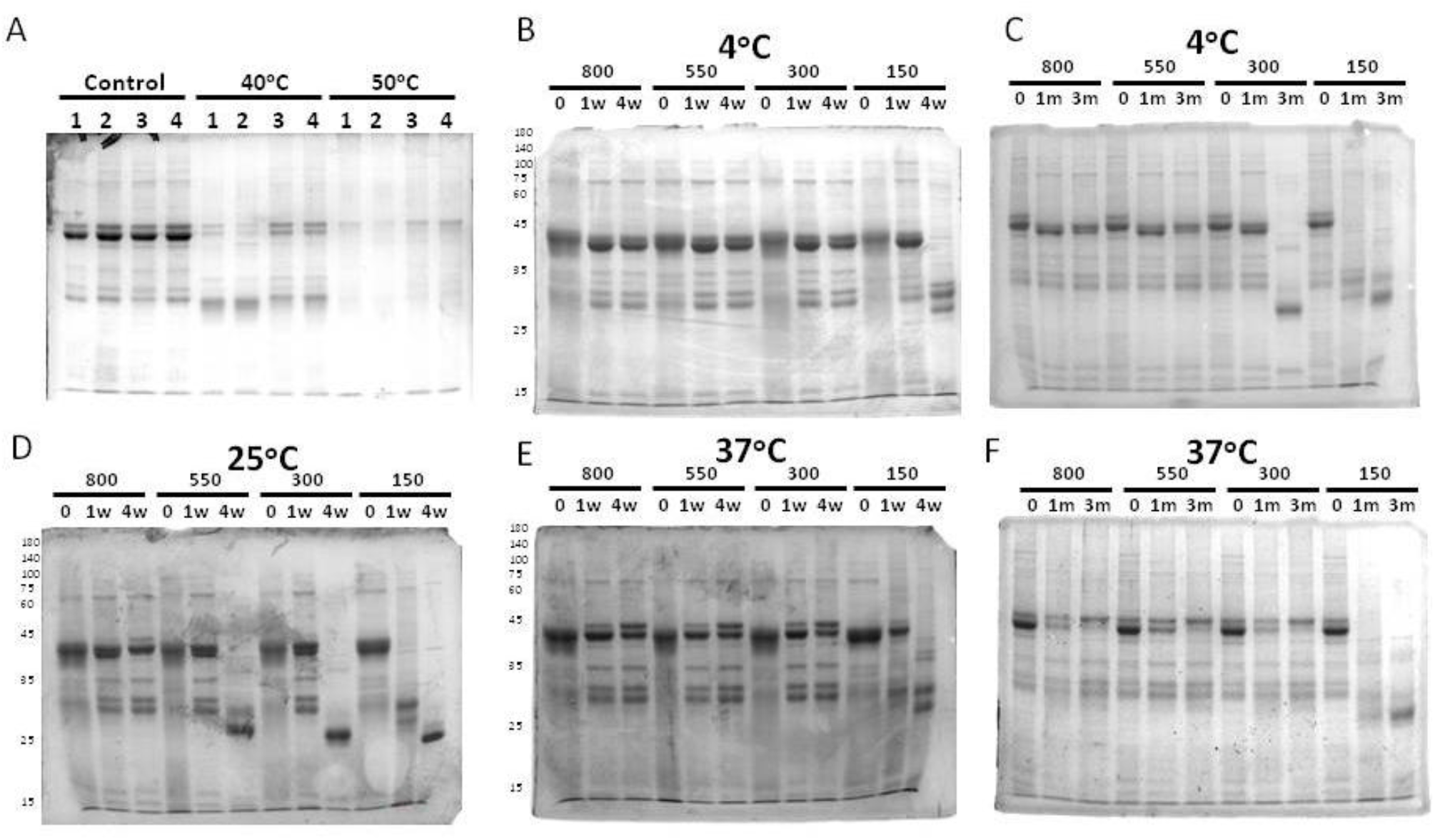
The analysis of protein nanoparticle stability using SDS-PAGE. The protein nanoparticle stability was analyzed by both SDS-PAGE and DLS. (A) Protein samples of 1) AH3-sfGFP-2xhM2e, 2) LY-sfGFP-2xhM2e, 3) RRLE-sfGFP-2xhM2e or 4) LYRRLE-sfGFP-2xhM2e were incubated in either 40°C for 4 weeks days and analyzed in 12% SDS-PAGE for protein stability. LYRRLE-sfGFP-2xhM2e protein nanoparticle was first reconstituted into phosphate buffer containing 150mM NaCl and then the NaCl concentration was adjusted to either 300mM, 550mM or 800mM before been stored in 4°C (B, C), 25°C (D) or 37°C (E, F) the period of time indicated. Protein samples were removed and processed then stored before been analyzed by SDS-PAGE.

These results provide strong evidences that the protein nanoparticle assembled by LYRRLE peptide is salt dependent and has good stability in high temperature (37 °C). This salt dependent nanoparticle stabilization is consistent with our assumption that the hydrophobic interaction is the major force that mediates monomer-monomer and dimer-dimer interactions.

## Discussion

Vaccine as a tool to prevent infectious diseases is the most cost effective strategy. Especially for attenuated viral vaccines like Vaccinia, MMR or oral polio vaccine, they produce long lasting even life time protective immune responses but these attenuated viral vaccines took decades for development. Apparently, this strategy will not be able to timely develop a vaccine to ward off emerging global pandemic like Covid-19. Although the new vaccine technology like DNA vaccine, mRNA vaccine or adenovirus based vaccine that can quickly develop a subunit vaccine after the genomic information of pathogen become available, but the immune responses generated are often declined to base level within a year (Feldman et al., 2019; Li et al., 2016; Modjarrad et al., 2019; Zhu et al., 2016). This short lived immune response may expose vaccinated people to the risk of breakthrough infection and in need of booster shot. In this study, we have created a self-assembled protein nanoparticle composed of a gain-of-function mutant of AH3 peptide that enables temperature and salt dependent protein nanoparticle thermostability. Protein nanoparticle assembled by this gain-of-function mutant is able to stimulate long lasting humoral immune response that is correlated with the thermostability of gain-of-function mutant. The surprising continued increase of anti-sfGFP IgG titer at 3 months post immunization is distinct from anti-hM2e IgG titer. This result suggests the continued presence of functional sfGFP protein without inserted hM2e peptide in immunized mice for 3 to 6 months. Whether these sfGFP is presented to immune system as loop hydrolyzed nanoparticle or single processed sfGFP is unknown. The increase of hydrophobic interaction by NaCl is due to the binding and removal of water molecules from protein in the presence of high NaCl concentration. Although the physiological NaCl concentration is maintained at 150mM, but there are other electrolytes in the body fluid that can serve the same role without disturbing electrical potential across cell membrane. So the in vitro stability results shown in this study may not represent what happened in injection site during immunization. The exact mechanism of this long lasting immune response is not known but it may be mediated by the presence of thermal stable protein nanoparticle that remains intact in injection site and stimulates the expression of long lasting plasma cell through continuous exposure, the same mechanism that accounts for the lifelong protection of attenuated viral vaccines (Amanna et al., 2007).

Fluorescent protein (FP) family is a group of proteins with a conserved sturdy barrel structure that encloses a fluorophore. Since the function of this beta strand constituted barrel is to provide a tightly controlled environment for fluorophore maturation and function, the thermostability of FP barrel can be improved by directed selection against fluorescent for mutants with high tolerance to structure destabilize (Tsien, 1998). The variability and stability of FPs make it a powerful tool as a vaccine carrier. First of all, the thermostability of FP partly contributes to the thermostability of LYRRLE-FP based protein nanoparticle, a desired vaccine property for achieving Global Vaccination Initiative. Second, because the diverse origins of FP (Dove et al., 2001), the LYRRLE-FP nanoparticle platform may be expanded into a line vaccine carriers that share a common format but each with distinct serum types. This feature of LYRRLE-FP platform may avoid the effects of pre-existing carrier antibody when LYRRLE-FP carrier is used in several vaccines. Third, the extensive studies on the structure of FPs and their applications have provided useful tools to incorporate heterologous antigens onto LYRRLE-FP based protein nanoparticle. With the features mentioned above, LYRRLE-FP format is suitable for the commercialization of a nanoparticle based subunit vaccine.

In this study, AH3 peptide derived from M2 protein of type A influenza strain H5N1 is found to induce nanoparticle formation when it was fused with a GFP. Previous studies about M2 amphipathic helical peptide focus on its roles in virus budding and proton pump anchorage. It mediates virus budding by generating membrane curvature by embedding its hydrophobic face in membrane bilayer (Roberts et al., 2013). And it anchors M2 ectopic domain on the viral envelope that serves as a proton pump that trigger envelope membrane fusion during infection (Hu et al., 2020). Here we discover a novel application of AH3, it serves as a nucleating center for protein nanoparticle assembly. This novel application is restricted to conditions when an AH3 peptide is fused with a fast folding protein like fluorescent protein. Fusing amphipathic helical peptide to other slow folding proteins tested causes misfolding of fusion protein apparently due to the aggregation of slow folding fusion proteins due to the presence of AH3 (unpublished results). So, the fast folding ability of sfGFP is the key to AH3-FP protein nanoparticle formation. Through protein modeling, we go further to identify two point mutations in AH3 peptide, I9L and K14E, that act synergistically to stabilize protein nanoparticle. The I9L mutation increases monomer-monomer intermolecular interaction, and the K14E mutation enables hydrogen bond formation that holds the tandem dimers together and enables removal of hydrophobic patch from surface. The synergistic effect of these two AH3 mutations contributes to a more stable protein nanoparticle and restricted it from binding cellular membrane none-specifically likely result in extending the lifetime of antibody responses.

The stabilization of LYRRLE-FP nanoparticle by high NaCl concentration is consistent with the assumption during structure modeling. Hypertonic saline up to 5.4% (6 folds concentration of normal saline, 0.9%) is routinely used in clinical trials involving study of neuromuscular pain. Intramuscular injection of hypertonic saline into muscle induces intense but fast tampered muscular pain with no detectable damage to injection site tissue afterward (Svendsen et al., 2005). The use of hypertonic saline (3.6% NaCl) in aluminum hydroxide formulated antigen was able to stimulate higher cellular and humoral immune responses than normal saline in mice (Luo et al., 2017). Other electrolytes or ingredients like sorbitol, sucrose, histidine or recombinant human serum albumin maybe added to decrease the use of NaCl to minimize interfering membrane potential post immunization. These electrolytes can distract water molecules from solvating proteins and keep protein nanoparticle bound by hydrophobic interaction intact. The fact that hypertonic saline is safe for human administration and has a potential benefit of further boosting immune responses is encouraging to apply LYRRLE-FP protein nanoparticle for the benefits of both stimulating long lasting immune responses and high thermostability during storage.

