## Supplemental for "A thermally stable protein nanoparticle that stimulates long lasting humoral immune response"

**Supplementary material**

**Figure Legend**

**Figure S1.**

Anti-GFP antibody titers of mice immunized by fusion proteins composed of amphipathic helical peptide and GFP. Mice were immunized twice with 14 days apart by intramuscular injection of 20 μg of fusion protein indicated. Sera collected at day 14 to day 16 post booster immunization were used for ELISA analysis. Data were compiled from 3 different individual experiments under the same experimental procedure. The ELISA plates were coated with 10μg/ml GFP and the 2$^{nd}$ antibody was used at a concentration of 1:2000 dilution. The other procedures were executed as standard protocol. (N=4 or 5) The result suggested the fusion of amphipathic helical peptide onto GFP increases the antigenicity of GFP by 2~3 log.

**Figure S2.**

Examining the protein stability of AH1-GFP and AH3-GFP fusion protein in ambient temperature. AH1-GFP and AH3-GFP transformed into BL21(DE3) or ClearColi competent cells for protein expression were induced at 15$^{o}$C overnight and then following the standard protocol for protein purification. Proteins were dialyzed into 1xPBS in a dialysis membrane with MWCO of 3000 Da overnight. (A) Purified proteins were kept in 37°C for one week or one month and then analyzed by SDS-PAGE and Coomassie blue staining. (B) Fusion proteins from ClearColi were kept in 37°C one month or RT (25°C) for one week or 5 months and then analyzed by SDS-PAGE and Coomassie blue staining. Results from figure S2A indicates AH1-GFP fusion protein stability in 37°C is reduced in the presence of LPS, but not AH3-GFP. Also, from Figure S2B, the result indicates the AH3-GFP is stable in both 25 °C and 37°C up to 5 months. But for AH1-GFP fusion protein, more than 70% of the protein was degraded into GFP in a week and completely converted to GFP in 5 months in 25 °C. The results suggest AH3-GFP is stable in both ambient temperatures tested.

Figure S3.

The hydrophobicity of the AH3-GFP protein nanoparticle was analyzed using density gradient ultracentrifugation. Protein samples were loaded on top of sucrose step gradients and then centrifuged at 35K rpm for 2 hrs in a SW41-Ti rotor. After centrifugation, fluorescent protein distribution was detected under 450nm LED light and pictures taken for analysis by ImageJ from NIH. (A) The centrifuge tube was loaded first with 1ml 45% (w/v) sucrose solution and then followed with 9ml of 15% (w/v) sucrose solution. One milliliter soluble fraction from each sample was loaded on top of the centrifuge tube before ultracentrifugation as described. Results of two samples were shown in figure S3B: GFP and AH3-sfGFP-2xhM2e. The results showed the GFP protein alone remained in the top of the centrifuge tube, but AH3-sfGFP-2xhM2e had been sedimented to the junction between 15%/45% sucrose solutions.

Supplementary materials

Figure S1

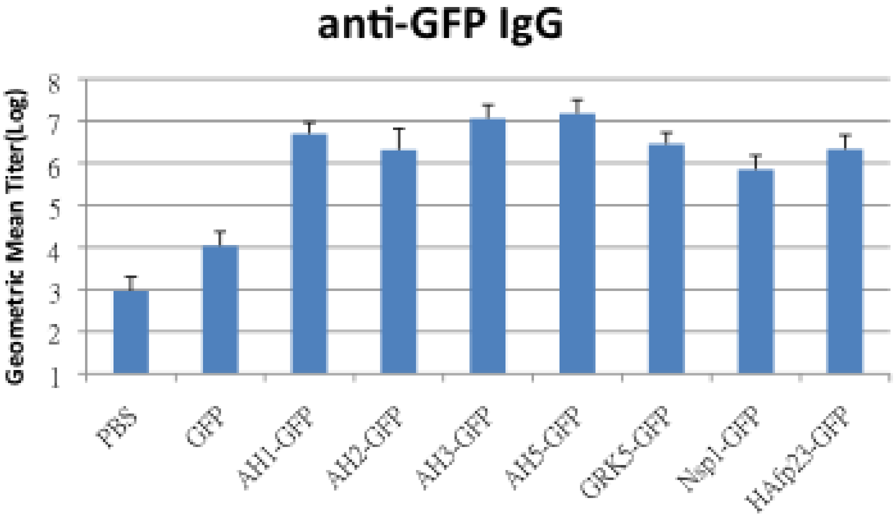


Figure S2

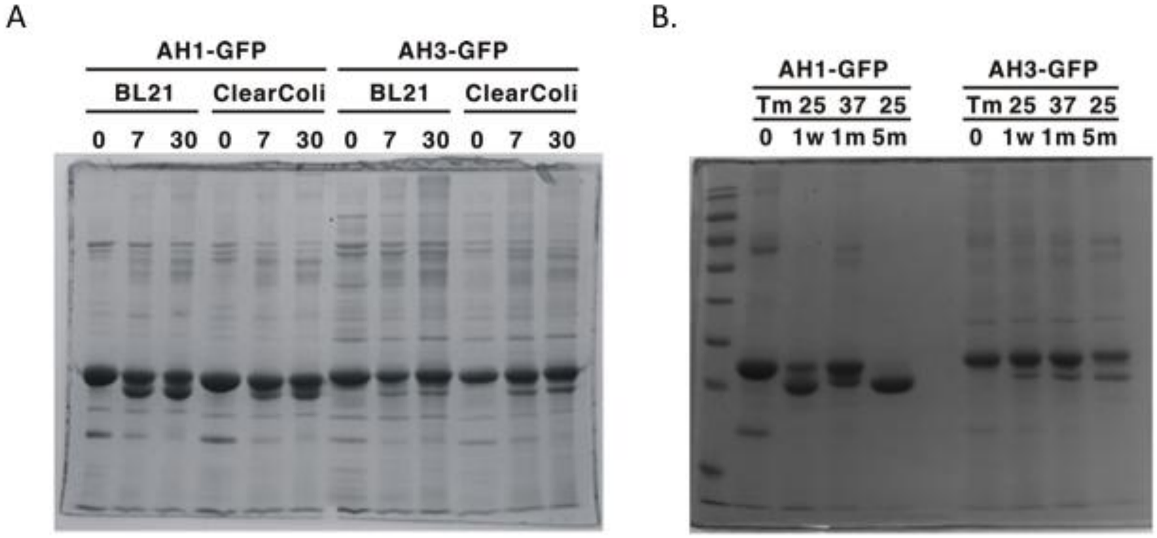

Figure S3

A

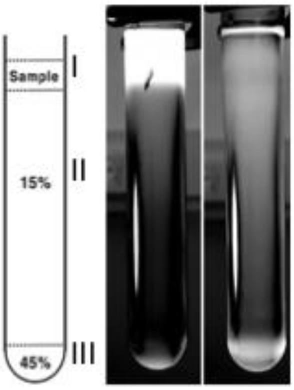

### Materials and Methods

#### Peptide information, expression and purification of recombinant protein

The AH3 peptide sequence is DRLFFKCIYRRLKYGLKRG. The sequence of peptide for anti-hM2e antibody titer ELISA is SLLTEVETPIRNEWGSRSNGSSDC. The peptide inserted in the insertion site of sfGFP is SLLTEVETPIRNEWGSRSN-GSSDSSGGSLLTEVETPIRNEWGSRSNGSSD. The protein expression vectors encoding target proteins were transformed into E coli competent cells using a heat shock transformation. Colonies of transformed bacteria with the desired vector were scraped from plate and inoculated in LB culture with an antibiotic. The bacterial culture was then growing exponentiallyd to OD600 between 0.5~0.7 before cooling down on an ice bath and protein expression was induced by 1mM IPTG at 20 $^{o}$C for 14-16 hours with 250 rpm shaking. After protein induction, bacteria were harvested by 5000 rpm centrifugation in a Sorvall SLC3000 rotor for 10 minutes. Bacterial pellet from 400 ml LB culture was re-suspended in 40 ml lysis buffer contains 10 mM Imidazole in 1XGF buffer (20 mM $Na(PO_4)$ pH7.4 and 300mM NaCl) for sonication. For Ni-NTA resin purification: 10 mM Imidazol was added in 1XGF buffer for bacteria lysis (Lysis buffer), 20 mM Imidazol was added in 1XGF buffer for column wash (Wash buffer), 500 mM Imidazol was added in 1XGF buffer for protein elution (Elution buffer). Bacteria were lysed using an ultrasonic sonicator (Misonix 3000) at 10 second on/20 second off cycles for 5 minutes at output level 5 in icy water. Insoluble cell debris was removed by centrifugation in 10000 rpm for 10 minutes using a Sorval SS34 rotor at 4 $^{o}$C. Soluble fraction containing the target protein was then used for purification by Ni-NTA resin as described in the user manual or been used for sedimentation ultracentrifugation. Purified proteins were stored in elution buffer in cold room before further processing. Before immunization or thermostability test, protein buffer was changed into 0.5x GF buffer using Sephadex-25 resin (GE,

### Acknowledgement

### Author contributions

Gunn-Guang Liou processed the protein nanoparticle samples and taking the TEM images. Ten-Tsao Wong designed and synthesized the pET27a based AH3-sfGFP expression vector. Ming-Chung Kan has done all the other works.

### Institutional review board statement

The mice immunization protocol had been reviewed and approved by IACUC of Fu Jen Catholic University

### Data availability statement

**Protein model can be accessed from Modelarchive.org**

### Conflicts of interest

**There is no conflict of interest.**
